# A Microphysiologic Model of the Cervical Epithelium Recapitulates Microbial, Immunologic, and Pathogenic Properties of Sexually Transmitted Infections

**DOI:** 10.1101/2025.07.21.665989

**Authors:** Katherine M. Nelson, Daniel J. Minahan, Vonetta L. Edwards, Ian J. Glomski, David J. Delgado Diaz, Keena Thomas, Forrest C. Walker, Patrik M. Bavoil, Isabelle Derré, Alison K. Criss, Jacques Ravel, Jason P. Gleghorn

## Abstract

Sexually transmitted infections (STIs) of the cervicovaginal mucosa are among the most common global infections. Clinical studies have revealed that susceptibility to STIs and the subsequent host responses they elicit are frequently associated with vaginal microbiota compositions that facilitate infection. Current monolayer cell culture and animal models fail to reproduce the multilevel complexity required to investigate these relationships simultaneously and/or with sufficient physiological relevance. To address this limitation, we have developed a microphysiologic system (MPS) that models human cervical tissue, its microbiota, and is susceptible to infection by two prominent genital pathogens, Chlamydia trachomatis and Neisseria gonorrhoeae. Significantly, this MPS platform recapitulates essential dynamic, polymicrobial, immune, and pathogenic features of chlamydial and gonococcal infections as they occur in humans. The low-cost MPS device requires no specialized equipment or specific expertise and was experimentally validated for both chlamydial and gonococcal infections across multiple non-engineering, remotely located laboratories, demonstrating its transferability and reproducibility. The MPS platform described herein provides a novel tool for expanded research into genital infections in a reconstituted system that closely mimics the cervical epithelium, a significant advance over existing models.

## Introduction

Sexually transmitted infections (STIs) of the cervicovaginal mucosa are among the most prevalent infectious diseases globally and result in multibillion-dollar economic losses annually (1, 2). In 2020, the World Health Organization reported more than one million new cases of chlamydia, gonorrhea, trichomoniasis, or syphilis each day among individuals aged 15-49 (3). Infection by *Chlamydia trachomatis* (*Ct*) and *Neisseria gonorrhoeae* (*Ng*), the causative agents of chlamydia and gonorrhea, respectively, are frequently associated with disruptions in the cervicovaginal microbiome(4, 5). The cervicovaginal microbiota, known to play a role in both health and disease(4, 6–10), can shift from an optimal Lactobacillus-dominated state to a nonoptimal dysbiotic state comprising diverse anaerobic bacterial species. This shift has been implicated in increased STI risk, and mucosal inflammation(5, 11–15).

The complex interactions among host genital epithelial cells, systemic and local immune defenses, the resident microbiota, and sexually transmitted pathogens remain poorly understood. While significant progress has been made by investigating these interactions in relative isolation using monolayer cell culture model systems, such models offer limited physiological relevance due to their inability to replicate the complexity of human tissues. More recently, cell culture insert models (*e*.*g*., Transwell^®^ systems) have been established to better understand the interactions between the host, the microbiota, and pathogens(16–20). However, these models have key limitations, including the lack of fluid flow, which precludes the study of the circulating immune responses, an essential aspect of host-pathogen interactions. Animal models, such as mice and other rodents(21–29), may more closely mimic certain aspects of natural infections in humans, but they are often poor surrogates. This is due, in part, to the host specificity of many STI pathogens and fundamental physiological differences between rodents and humans. For example, the immune response in mice differs substantially from that in humans; essential receptor proteins and nutrient sources required by the pathogens to establish an infection may be absent, and the murine cervicovaginal microbiota is significantly different from its human counterpart(30–34). A major barrier to mechanistic insight has been the lack of a physiologically relevant, experimentally accessible human tissue model that captures the dynamic and multifactorial nature of the cervical microenvironment. Such a model is critical for advancing our understanding of the immunologic, microbial, and pathogenic determinants of STIs susceptibility and disease progression in a clinically representative environment.

The fabrication of microphysiological system (MPS) devices and the co-culture of multiple mammalian cell types within these devices are well-established, albeit specialized, in the field of biomedical engineering(35–40). Commercially available MPS platforms now exist to model various human organ systems. However, their fabrication and use often require access to specialized instruments, facilities, and technical expertise, making them impractical for routine use in laboratories without engineering infrastructure. Here, we describe the design, biological validation, and successful multi-laboratory implementation of a modular cervical MPS device across several independent, non-engineering laboratories. To accomplish this, we assembled an interdisciplinary team combining the expertise of biomedical engineers with that of researchers specializing in the microbiome, *Ct*, and *Ng*. Together, we designed, validated, and ensured the reproducibility of a physiologically relevant cervical model. We demonstrate that this model supports the complete developmental cycle of *Ct* infection, microbiota-mediated protection against *Ct* infection, and neutrophil recruitment and transepithelial migration in response to *Ng* infection. This MPS provides a new and versatile platform for mechanistic studies of host-microbiota-pathogen interactions in the female reproductive tract and offers a broadly accessible tool to studying STI pathogenesis, microbial therapeutics, and mucosal immunity. Ultimately, this low-cost, reproducible platform, with accompanying protocols, is designed to be easily adopted with minimal training by non-engineering research laboratories, thereby lowering the technical barrier for advanced experimental modeling in reproductive health research.

## Methods

### Modular MPS design and fabrication

Devices were fabricated as described previously(41). Briefly, insert geometries were designed in SolidWorks and cut into silicone sheets with pressure-sensitive adhesive via a vinyl cutter (USCutter). A 3 µm-pore polycarbonate membrane (Cytiva, 10418306) was placed between two silicone layers to form the insert. The cassette lid and base were fabricated from acrylic sheets and laser cut (Full Spectrum Laser, Hobby Series 20×12) to the proper geometry. The top acrylic piece was raster etched to create a recess for a round coverslip, which was inserted and secured with adhesive.

### Human cervical cell cultures in the MPS insert

#### Cell types and maintenance culture

A2EN cervical epithelial cells were a gift from Dr. Alison J. Quayle (Louisiana State University Health Sciences Center)(42). A2EN cells were cultured in EpiLife® phenol red-free medium (Gibco MEPICFPRF 500) supplemented with 60 *µ*M CaCl2, EpiLife® Defined Growth Supplement (EDGS; Gibco S0125), and L-glutamine (Gibco 35050-0610). BJ fibroblasts (ATCC CRL-2522) were cultured in Dulbecco’s Modified Eagle Medium (DMEM; Gibco 11965-092) supplemented with 10% heat-inactivated fetal bovine serum (FBS). Primary cervical epithelial cells (HCxEC; LifeLine Cell Technologies FC0080) were cultured in ReproLife CX Medium Complete Kit (ReproLifeTM CX Basal Medium and ReproLifeTM CX LifeFactors Kit, LL-0072). HeLa cells (ATCC CCL2) were cultured in Dulbecco’s Modified Eagle Medium (DMEM; Gibco 11965-092) supplemented with 10% heat-inactivated fetal bovine serum (FBS). All cells were propagated at 37^°^C 5% CO_2_. Penicillin and streptomycin (Gibco 15140-122) were added to culture media when cells were seeded on the device but were washed and removed at least one day before the addition of bacteria.

#### Cell cultures in MPS device

For static culture, inserts were placed in a custom-fabricated well (Protolabs) designed to hold medium and provide access to both the basal and apical compartments. While seated in the well, the polycarbonate membrane of the insert was coated on both sides with 0.5 mg/mL fibronectin (Millipore FC010) for 45 minutes at 37°C. Excess fibronectin was then removed using a pipette. BJ fibroblasts were seeded on the upright basal side of the polycarbonate membrane at a density of 2×10^5^ cells/insert and allowed to adhere for at least 2-4 hours at 37^°^C 5% CO_2_. After this initial adhesion period, DMEM supplemented with 10% FBS medium was added to the well, and the insert was inverted to position BJ fibroblasts in contact with the medium. Then A2EN or HCxEC cervical epithelial cells were seeded on the now upright apical side of the polycarbonate membrane at a seeding density of 2×10^5^ cells/insert in epithelial cell medium. Cells were allowed to adhere for 2-14 hours, under hypoxic conditions (37°C, 5% CO_2_, 3% O_2_); less adherence time was required for A2EN cells, while longer adherence times were required for HCxEC. After incubation, both the basal and apical media were removed. The basal medium was replenished with the appropriate epithelial cell medium, which was changed every 48 h, and no fresh apical medium was added. Inserts were maintained under static culture conditions until confluent BJ fibroblast and epithelial cells monolayers formed, and an air-liquid interface (ALI) was established (37°C, 5% CO_2_, 3% O_2_) (approximately 7-13 days, depending on the epithelial cell types).

#### Fluidic culture set-up and flow conditions

For dynamic culture conditions, once the cervical epithelial and fibroblast monolayers were established, the insert was clamped into the acrylic cassette and connected to a peristaltic pump via tubbing. Flow was then initiated, as previously described(41). Medium flow was maintained at 2.5-5 µL/min through both the apical (epithelial) and basal channels (37°C, 5% CO_2_, 3% O_2_).

### Characterization of cervical cell culture in the MPS device

#### Assessment of barrier formation

To assess the formation of a complete epithelial monolayer, barrier function was evaluated using a FITC-dextran diffusion assay. Medium containing 0.1 mg/mL 150 kDa FITC-dextran was added to the apical side of fibronectin-coated inserts, either cellfree or containing a co-culture of monolayers of A2EN epithelial cells and BJ fibroblasts. At designated timepoints, medium samples were collected from the basal compartment, and fluorescence was measured using a fluorometer (excitation: 488 nm; emission: 517 nm). Fluorescence values were compared to a standard curve to quantify the amount of FITC-dextran that had diffused across the insert membrane and cell layers(43). Statistical analysis was performed using an unpaired t-test in GraphPad Prism software.

#### Immunofluorescent staining

Immunofluorescent staining was used to visualize epithelial and fibroblast monolayers. Inserts were removed from the wells (static) or cassette (flow), rinsed with PBS, and fixed with 4% paraformaldehyde (PFA). Following fixation, cells were permeabilized with 0.1% Triton X-100 in PBS and blocked with 1-3% bovine serum albumin (BSA). Cells were incubated overnight at 4°C with primary antibodies against mucin 5B (Sigma, HPA008246 at 1:20), E-cadherin (BD transduction laboratories, 610181 at 1:200), and/or occludin (Invitrogen, 40-4700; diluted 1:100). Alexa Fluor-conjugated secondary antibodies (Molecular Probes, A11037; diluted1:500 or Invitrogen, A11029, A11008, A11030, A11038; each diluted 1:200) were added and incubated for 2 hours at room temperature in the dark to visualize the primary antibodies. F-actin was stained with Phalloidin (Invitrogen, A12379, A22283; diluted 1:400) for 2 hours at room temperature in the dark. Nuclear straining was performed using either SYTOX-green (Invitrogen, S7020; diluted 1:50,000) or Hoechst (Invitrogen, H3570 at 1:500) at room temperature in the dark. Following staining, the membrane was removed from the insert by peeling and mounted onto glass slides with coverslips using VectaShield (Vector Laboratories, H1000) and stored at 4°C in the dark until imaging.

#### Imaging of fixed insert

Fixed inserts were imaged using either a Leica DMi8 spinning disc confocal microscope equipped with a dry long-working distance 63X objective, an Andor iXon ULTRA 888BV EMCCD camera, and a CSU-W1 confocal scanner unit or a Nikon A1 Laser confocal microscope with a 60X objective. For 2D imaging, a 20X objective on an EVOS FL Auto 5000 imaging system (ThermoFisher) was used. To visualize the full insert channel, multiple fields of view were captured using a 4X objective on a Nikon Eclipse TE200-U microscope. These images were manually stitched and merged in PowerPoint. Image processing and analysis were performed using Imaris software (Bitplane, Belfast, United Kingdom) or FIJI/Image-J software (NIH, Bethesda, United States).

### Bacterial strains and microbiota consortia

Reconstructed microbiota consortia were generated by combining selected clinical isolates to represent distinct cervicovaginal community types. The ‘optimal’ microbiota consortium consisted of three clinical strains of *Lactobacillus crispatus* (*Lc1*: C0241A1, *Lc2*: C0231A1, *Lc3*: C0103E5), while the ‘non-optimal’ microbiota consortium was composed of two *Gardnerella vaginalis* strains (*Gv1*: C0179B4, *Gv2*: C101A1) and one *Prevotella amnii* strain (*Pa1*: C0088E5), anaerobic bacteria typically found during episodes of bacterial vaginosis. Clinical isolates were initially cultured individually in S-broth under anaerobic conditions until they reached log phase (6 hours for *L. crispatus* strains; 24 hours for *Gv1, Gv2*, and *Pa1*). Cultures were quantified by measuring optical density at 600 nm (OD600), and strains within each consortium were mixed in a 1:1:1 ratio to create a final volume of 3 mL (1×10^8^ bacteria/mL). These mixtures were aliquoted in 500 µL and stored at −80°C as stock in S-broth containing 10% glycerol. *Chlamydia trachomatis* (*Ct*) lymphogranuloma venereum (LGV) serovar L2 (L2/434/Bu ATCC VR-902B) was used in the study. Plaque-purified *Ct* strains constitutively expressing red fluorescent protein mCherry (mCh *Ct*) or the reticulate-to-elementary body transition reporter (RB to EB *Ct*), were previously described(44, 45). Propagation assays for *Ct* were performed according to established protocols(46).

A piliated derivative of *Neisseria gonorrhoeae* (*Ng*) strain FA1090, constitutively expressing OpaD(47) and carrying a plasmid that encodes green fluorescent protein (GFP)(48, 49), was used to infect the epithelial models.

### Microbiota consortia addition and characterization in the MPS device

Prior to inoculation, one loopful of frozen optimal and non-optimal consortia was inoculated in 1 mL of S-Broth medium, grown anaerobically without shaking for 6 hours, and OD600 was measured to determine bacterial concentration according to 1OD = 1×10^8^ bacteria. The volume required for 4×10^5^/mL was added to 1 mL of HBSS++ (Hank’s Balanced Salt Solution containing 1.8 g/L glucose and 3.4 g/L maltose), and 50 µL of this suspension (containing 2×10^4^ bacteria) was added to each insert containing epithelial cells and fibroblast monolayers. The inserts were then incubated under hypoxic conditions (5% CO_2_, 3% O_2_) for 48 hours. An aliquot of the inoculum (0h) was stored at −20°C for quantification of each strain by qPCR. After 48 hours, the inserts were removed and disassembled with forceps under sterile conditions in a biosafety cabinet. The membrane was collected and cut with a sterile scalpel and combined with any medium from the apical channel into an Eppendorf tube containing 100 µL deionized water and stored at −20°C. Bacterial DNA was extracted using the AllPrep PowerViral DNA Kit (Qiagen, 28000-50) with the lysis step performed directly in the Eppendorf tube containing the membrane. A multiplex strain-specific qPCR assay was designed and validated for each consortium, allowing for the quantification of each strain genome copy number within the consortia. Primer and probe sequences for these assays are listed in **Supplementary Table S1**. Data analysis and graphing were performed using GraphPad Prism software. Statistical significance was assessed using ANOVA and Student t-tests.

#### Effect of consortia on the cervical epithelium in the MPS device

Cervical epithelial cells (A2EN and HCxEC) were cultured on inserts under static conditions until polarization was achieved. Optimal and non-optimal consortia (2×10^4^ bacteria in HBSS++) were added to each insert and incubated under hypoxic conditions (5% CO, 3% O) for 48 hours. To assess changes in cell viability, epithelial cells on the inserts were stained using the Biotium Cell Viability/Cytotoxicity kit (Biotium; 30002 at 1:1000) and imaged. Basal medium was collected after incubation, stored at −80°C, and later analyzed by ELISA to evaluate host epithelial immune response. Specifically, inflammatory cytokines IL-8 and IL-1*β* basal levels were measured using Quantikine ELISA kits (Invitrogen D8000C for IL-8 and DLB50 for IL-1*β*) following the manufacturer’s recommendations. Data visualization and statistical analysis were performed in GraphPad Prism software. Statistical significance was assessed using ANOVA.

### *Ct* infection and analysis in the cervical MPS device

Following co-culture of A2EN epithelial cells and BJ fibroblasts, the culture medium in each insert apical compartment was replaced with phenol red-free DMEM (Gibco, 31053-028) supplemented with 2% charcoal-stripped, heat-inactivated fetal bovine serum (FBS; Gibco, 12676029) and 1X GlutaMAX (Gibco, 35050-061) to support productive *Ct* infection. *Ct* strains were diluted in the same phenol red-free DMEM supplemented with 2% charcoal-stripped, heat-inactivated FBS to achieve a multiplicity of infection (MOI) of 0.5. A 40 *µ*L of the *Ct* suspension was added to the apical surface of the A2EN epithelial layer, and inserts were incubated at 37°C under hypoxic conditions (5% CO, 3% O) for the indicated times.

#### Live imaging of Ct-infected cervical epithelial cells on the inserts

To monitor *Ct* inclusion formation and growth over time, inserts were carefully transferred to a 100 mm tissue culture dish with the epithelial side facing downward and maintained in medium to prevent desiccation. Live imaging was performed every 24 hours using a Nikon Eclipse TE200-U epifluorescence microscope. Images were processed using NIS-Elements AR 6.02.03 (Nikon Instruments, Tokyo, Japan).

#### Staining and imaging of Ct-infected cervical epithelial cells on the inserts

Inserts were carefully transferred to a fresh 100 mm tissue culture dish, and cells in both apical and basal channels were fixed with 4% paraformaldehyde (PFA) for 10 minutes, followed by three PBS rinses. After fixation, inserts were transferred to a one-chamber coverglass system (Lab-Tek II, 155360), and channels were filled with DABCO antifade-containing medium. Imaging was performed using 4X and 10X objectives on a Nikon Eclipse TE200-U epifluorescence microscope. Images were processed using NIS-Elements AR 6.02.03 (Nikon Instruments, Tokyo, Japan).

#### Measurements of Ct inclusion size and reticulate body (RB) to elementary body (EB) conversion

To quantify inclusion size, the surface area of each *Ct* inclusion within a field of view was determined using the “Measure Objects” function of NIS-Elements software in the mCherry fluorescence channel. For analysis of RB to EB conversion, a similar approach was applied using the mCherry and GFP fluorescence channels. Three fields of view were analyzed per insert at each time point. Data were graphed and analyzed using GraphPad Prism software.

### Primary and secondary *Ct* infection on primary cervical cells post-exposure to microbiota consortia

Primary cervical epithelial cells (HCxEC) were cultured on inserts in ReproLife CX Complete Medium for 10 days under static conditions until polarization was achieved. Either 2×10^4^ of the optimal or the non-optimal microbiota consortia were inoculated in HBSS++ medium on each insert and incubated under hypoxic conditions (5% CO, 3% O) for 48 hours. *Ct* serovar L2 was added to the HBSS++ on the apical epithelial cell surface at an MOI of 1 and incubated for 2 hours at room temperature with gentle rocking. Following infection, epithelial cells were rinsed with PBS and incubated under hypoxic conditions for an additional 45 hours. Following incubation, epithelial cells were rinsed, fixed in 4% PFA, and permeabilized with 0.1% Triton X-100. Chlamydial inclusions were stained using the D3DFA Chlamydiae Culture Confirmation Kit (Quidel Ortho, 01-040000 at 1:20) for 30 minutes at 37°C. Cells were imaged using a Nikon A1 confocal microscope, and the number of epithelial cell nuclei and chlamydial inclusions was quantified.

For secondary infection, the inserts were removed and disassembled with forceps under sterile conditions in a biosafety cabinet. The membrane was collected and cut with a sterile scalpel and placed into an Eppendorf tube containing 200 µL of water. After vortexing for 3 min, the *Ct*-infected HCxEC epithelial cells from the primary infection were lysed by passing them through a 25-gauge needle. A total of 800 µL of DMEM cell culture medium was added, and 250 µL of this lysate was added to duplicate wells in a 48-well plate containing HeLa cells, and rocked for 2 hours at room temperature, then rinsed with PBS. Fresh HeLa epithelial cell culture medium was added, and cells were incubated under hypoxic conditions (5% CO, 3% O) for an additional 22 hours. Following incubation, cells were stained for chlamydiae and epithelial cell nuclei, imaged using an EVOS FL Auto 5000 imaging system, and analyzed using CellProfiler software (Broad Institute, MA, United States). Data visualization and analyses were performed in GraphPad Prism software, and statistical analyses were performed using Student’s t-tests.

### *Ng* infection and analysis in the cervical MPS device

Five *Ng* colonies from an overnight culture of a frozen stock were streaked on GC medium base (Difco 228920) agar (Invitrogen 30391-023) plates containing Kellogg’s supplements(50) (0.4% (w/v) glucose, 0.01% (w/v) glutamine, 0.00002% (w/v) thiamine pyrophosphate, 0.00005% (w/v) Fe(NO_3_)_3_•9H_2_O(51) containing 0.5 µg/mL chloramphenicol to select for GFP expression. Plates were incubated for 18 hours to form a confluent bacterial lawn. Bacteria were harvested using a polyester-tipped applicator and resuspended into HBSS (Gibco 14025092) supplemented with 10 mM HEPES (pH 7.4) (Sigma H4034) and 5 mM NaHCO_3_ (Sigma S6014) (HHB). A total of 3×10^8^ CFU of *Ng* (based on 1×10^8^ CFU/0.16 OD550) were pelleted and resuspended in 1 mL of a 1:1000 dilution of CellTrace Far Red (CTFR; ThermoFisher C34572) in HHB. For static insert cultures, CTFR-labeled bacteria were pelleted, washed, and resuspended in epithelial cell medium to a concentration of 6.6×10^7^ CFU/mL. A 30 µL aliquot (2×10^6^ CFU) was added to the epithelial surface of each insert to achieve an MOI of 10. Inserts were incubated at 37 C in 5% CO_2_, 3% O_2_ for 1 hour(52). Following incubation, epithelial surfaces were gently washed with epithelial cell medium by pipetting and further incubated for the indicated durations. For infection under flow conditions, CTFR-labeled *Ng* were resuspended to a concentration of 4×10^7^ CFU/mL. A 50 µL aliquot (2×10^6^ CFU, or an MOI of 10) was injected into the inflow port of the flow cassette until bacterial turbidity was observed at the exit port. Flow was paused during the 1-hour infection period and restarted afterward by flushing fresh epithelial cell medium to remove bacteria from the apical channel. Flow was resumed for the indicated durations.

#### Assessment of Ng infection

At the indicated time points, inserts were removed and disassembled with forceps under sterile conditions in a biosafety cabinet. Epithelial cells were dissociated from the membrane with a 1% saponin solution in PBS. *Ng* CFUs were quantified by serial dilution of the lysates, plated on GCB agar, and incubated overnight. For imaging, inserts were fixed in 4% PFA, permeabilized with 1% Triton-X100 in PBS, and blocked with 2% BSA in PBS. F-actin was stained using Texas Red-phalloidin (ThermoFisher T7471) in 2% BSA PBS. Images from the intact insert were acquired with a dry long-working distance 63x objective and membranes from disassembled inserts were acquired with a 40x wet objective on a Zeiss LSM900 confocal microscope and processed using Zen Blue software.

### Neutrophil isolation, labeling, and transmigration assays

#### Immune cell isolation and purification

Human subjects research was approved by the University of Virginia Institutional Review Board for Health Sciences Research under protocol 13909, and informed written consent was obtained from all donors. Primary human polymorphonuclear neutrophils (PMNs) were isolated from peripheral blood of healthy donors(53) following the institutional Human Subjects in Research guidelines. Briefly, blood was collected in heparin-coated vacutainer tubes and subjected to dextran sedimentation to enrich leukocytes. Granulocytes were purified by Ficoll-PaqueTM density gradient centrifugation in DPBS without calcium or magnesium (Gibco 14190) supplemented with 0.1% glucose (DPBSG; Ricca Chemical). The granulocyte pellet was resuspended, erythrocytes were lysed with endotoxin-free water, and the remaining cells were washed and resuspended in DPBSG on ice. Cell counts were determined using a hemocytometer. PMNs were labeled with a 1:1,000 dilution of CellTrace Violet (ThermoFisher C34557) in DPBSG for 20 minutes at room temperature, washed and resuspended in epithelial cell medium.

#### Immune cell addition and migration

Inserts containing A2EN epithelial cells and BJ fibroblasts cultured under hypoxic conditions (5% CO_2_, 3% O_2_) without flow for 7 days were infected with *Ng* (MOI of 10) in the epithelial channel for 1 hour, as described above, and flipped over so that the epithelial surface was submerged downward in the well medium. As a positive control for PMN migration, 10 µM of the chemoattractant formylated-methionine-leucine-phenylalanine (fMLF, Sigma 47729)(16) was added to the epithelial surface. PMNs (1×10^6^) were added to the basal BJ fibroblast channel, and inserts were incubated under hypoxic conditions (5% CO_2_, 3% O_2_).

#### Characterization of PMN migration

At the indicated time points, basal medium from the well was collected to assess PMNs transmigration. Myeloperoxidase (MPO) activity was measured as a proxy for PMN abundance using a chromogenic substrate (ABTST Chromophore Diammonium Salt; EMD Millipore) as previously described(16). Readings were done on a Wallac Victor-2 1420 Multilabel Counter (Perkin-Elmer) with 1-second readings at 405 nm, and values were quantified against an MPO standard curve. For imaging, inserts were removed from culture at the indicated time points, fixed in 4% PFA, permeabilized with 1% Triton-X100 in PBS, blocked with 2% BSA in PBS, and F-actin stained with Texas Red-phalloidin (ThermoFisher T7471) in 2% BSA PBS. Images were acquired with a Zeiss LSM900 confocal microscope as indicated above.

### Technology transfer and cross-laboratory implementation

To evaluate reproducibility and user accessibility, standardized protocols for device fabrication, cervical cell seeding, microbial consortia inoculation, and infection workflows were developed and implemented across four independent laboratories. The protocols were disseminated as written guides and instructional videos. All participating laboratories successfully reproduced model establishment under static and flow conditions, and implemented microbiome, *Ct*, or *Ng* infections and downstream analyses as needed. These results demonstrated the robustness and transferability of the platform across research settings. For additional information, including access to protocol, devices, or opportunities for collaboration, please visit https://www.gleghornlab.com/resources

## Results

### Cooperative and iterative design, cross-laboratory validation, and reproducibility

We developed a modular MPS device using a removable insert and reusable cassette system that supports cell culture under static or flow conditions and is compatible with standard downstream biological assays. The device was specifically designed for fabrication without the need for cleanroom facilities, allowing use with conventional laboratory equipment and workflows. It was validated for compatibility with epithelial, bacterial, and immune components across multiple independent laboratories. To enable broad adoption, we assembled an interdisciplinary team of researchers to optimize technology transfer. Written protocols and instructional videos were co-developed to support this process (**Figure 1A**). The resulting MPS device was iteratively refined to meet bioengineering requirements by incorporating suggestions from end-users into the design, with the overarching goal of modeling host-pathogen-microbiome interactions in a biomimetic model of the human cervical epithelial environment. The tissue-specific device builds on a previously described modular MPS device platform developed by the Gleghorn Lab(41), incorporating a single-use removable insert for cell culture that can be clamped into a reusable cassette (**Supplemental Figure S1**). The insert-cassette configuration allowed for straightforward microfluidic setup and cell culture workflows (**Figure 1B**), using kits that fit easily within a standard tissue culture incubator (**Figure 1C**).

**Fig. 1.**
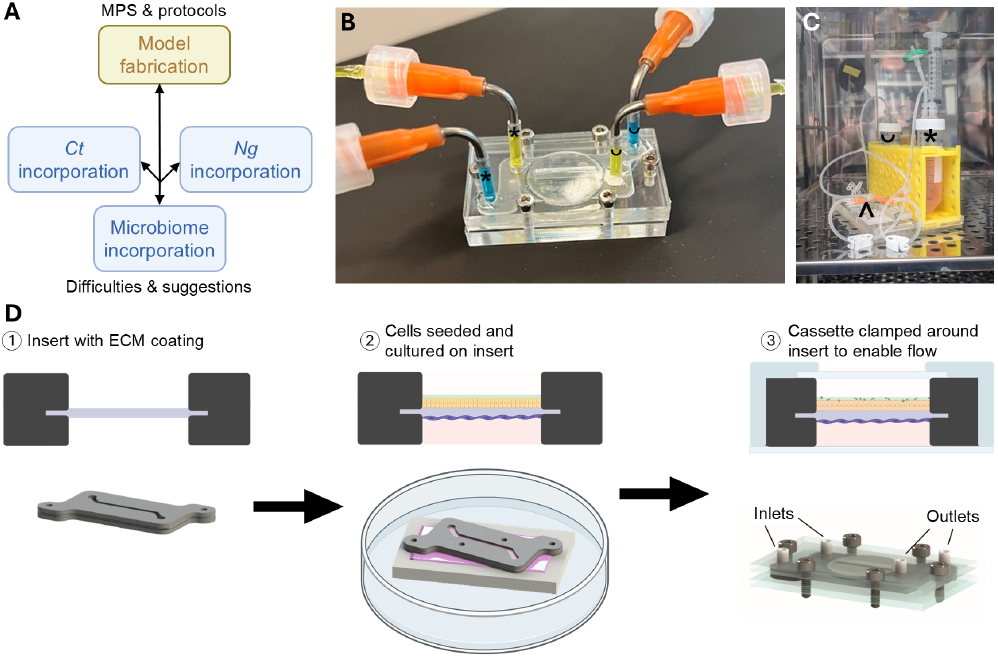
The microphysiologic system (MPS). **(A)** Flowchart of materials and information sharing between the collaborating labs. **(B)** Image of real device under flow with food dye showing the epithelial and basal channels and **C)** the full setup within the incubator. Inlet (*) and outlet (∪) reservoirs are held in yellow conical tube rack and connected to the device (^) via tubing at the inlets (*) and outlets (∪) in B. **(D)** Steps of cell culture in the insert outside of the device before clamping in the cassette to add fluid flow. 1 – Membrane (purple) in the insert (grey) is coated with ECM proteins to enable cell adherence. 2 – Cervical cells (yellow) and fibroblasts (dark purple) are seeded and cultured in media (pink) in the insert outside of the cassette to enable easier cell culture. 3 – Bacteria (green) are added, and the cassette (blue) is clamped around the insert to enable fluid flow.

To support cell adherence, the membrane in a fresh insert was coated with extracellular matrix (ECM) proteins, similar to a standard Transwell® insert. Fibroblasts and cervical epithelial cells were then seeded on opposite sides of the insert in a standard Petri dish. The absence of microfluidic channel constraints during cell seeding and monolayer formation enabled simplified operation by users without microfluidics expertise. After cell seeding, experiments could be conducted under static conditions or transitioned to dynamic flow by clamping the insert into the cassette (**Figure 1D**). The insert-cassette system was connected to medium reservoirs via tubing and a peristaltic pump to initiate controlled fluid flow across the monolayers in microfluidic channels (**Figure 1C**). This modular design lowers the technical barrier to adoption for non-engineering laboratories and facilitates broader use of physiologically relevant organ-on-chip models in infection biology research.

### Human cervical epithelial cells and fibroblasts form polarized, barrier-forming monolayers

A2EN cells, an immortalized cervical epithelial cell line commonly used to study STIs(54–56), were used to establish a cervical MPS model. A2EN cells were seeded on the apical (epithelial) side of the insert and feeder BJ fibroblasts were seeded on the basal side, cultured either under static conditions (**Figure 2A**) or exposed to shear stress via medium flow at 5 *µ*L/min for 7 days (**Figure 2B**). BJ fibroblasts were required for consistent cervical epithelial cell growth, as confirmed by visual inspection and experimental validation (**Supplemental Figure S2A**). A2EN cells formed confluent monolayers and maintained E-cadherin expression, indicative of stable epithelial junctions (**Figure 2C**). One of the advantages of the MPS system is its ability to support tissue-like architecture and integrate physiologically relevant parameters to cell culture settings. In the configuration of the device developed here, A2EN cervical cells in the apical (epithelial) channel and BJ fibroblasts in the basal channel recreated a stratified, tissue-like spatial organization (**Figure 2D**). To functionally assess barrier function of the epithelium, a FITC-dextran diffusion assay was performed (**Figure 2E**). Inserts containing both A2EN epithelial cells and BJ fibroblasts showed about a 5-fold decrease in FITC-dextran diffusion compared to cell-free inserts, indicating effective barrier formation by the co-cultured monolayers(43, 57). We next tested the compatibility of the MPS device with primary cervical cells. Using HCxEC primary cervical cells, we observed the formation of a confluent monolayer that produced mucin, a hallmark of differentiated primary cervical cells (**Figures 2FG**). Primary cervical cells (HCxEC) required a longer (11 days) culture period to reach confluency compared to the immortalized A2EN cervical cell line (6 days) (**Supplemental Figure S2B**). Together, these results demonstrated that the MPS device supports the formation of functional cervical epithelial models with immortalized and primary cells under static and flow (shear stress) conditions. Establishing a stable epithelial barrier is critical for studying not only infection dynamics but also directional signaling, microbial translocation, and immune cell recruitment across polarized epithelia.

**Fig. 2.**
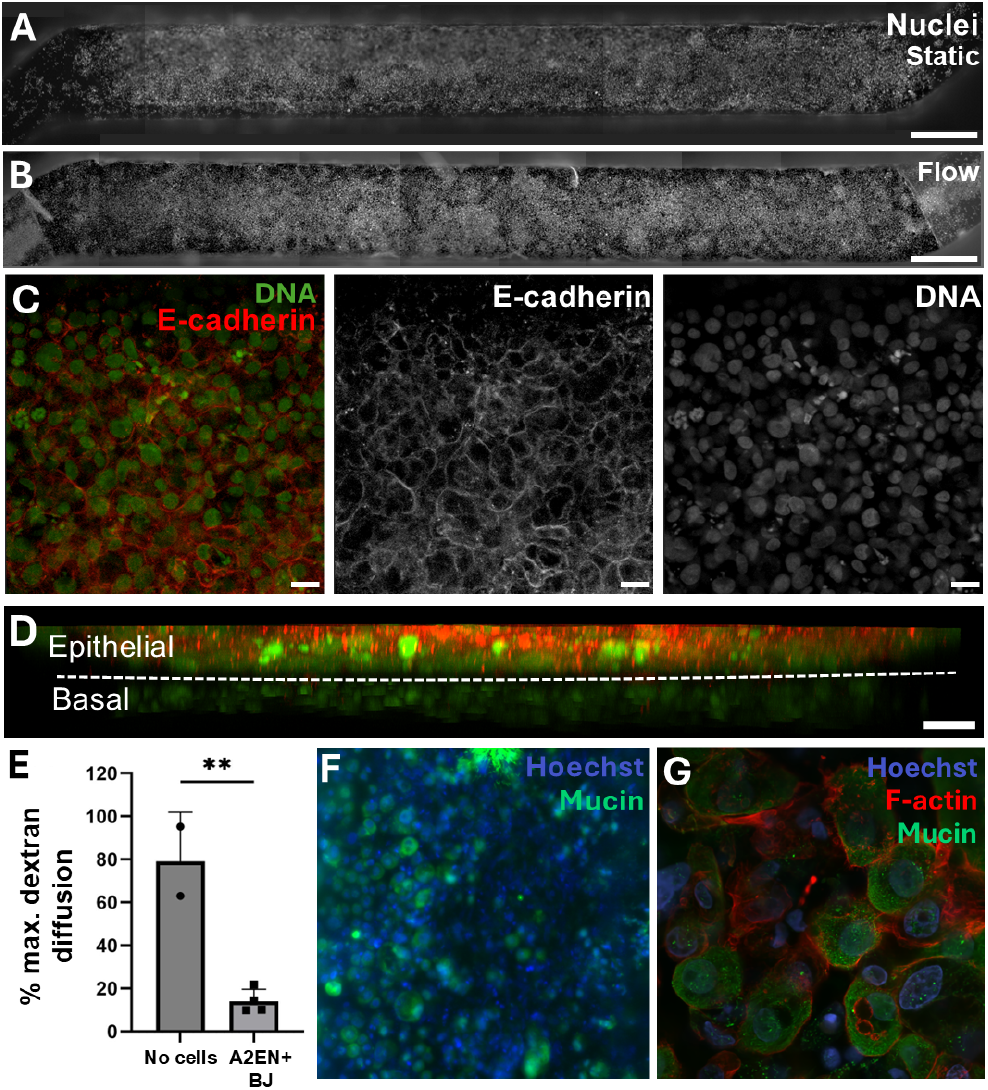
Establishment of a human cervical MPS incorporating cervical cells in the epithelial channel and BJ fibroblasts in the basal channel. **(A-B)** Epifluorescence microscopy top-view images of the cervical MPS epithelium with A2EN cells in the epithelial channel cultured under **(A)** static conditions or **(B)** continuous flow conditions at 5 µL/min for 7 days, stained with Hoechst DNA stain (white). Scale bar, 1 mm. **(C)** XY confocal images of a A2EN cervical epithelium stained for E-cadherin (red) and nuclei (SYTOX-Green, green), after 7 days of static culture. Merged corresponding monochrome split images shown. Scale bar, 20 µm. **(D)** XZ side-view as in C. Dashed line is the membrane. Scale bar, 20 µm **(E)** Functional barrier permeability assessment of an empty insert (no cells) and a fully confluent A2EN cervical epithelium co-cultured with BJ fibroblasts assessed using a 150 KDa FITC-dextran permeability assay Data were analyzed with an unpaired t-test. **(F-G)** Top-view of the epithelial channel with primary cervical cells (HCxEC) stained for mucin 5B (MUC5B, green), F-actin (red), and nuclei (Hoechst, blue) at day 11 of culture, under static conditions. Scale bars are 10 and 20 µm, respectively.

### Reconstructed optimal and non-optimal microbiota consortia grow stably and differentially modulate host responses

A2EN cervical epithelial cells or HCxEC primary cervical cells were co-cultured with BJ fibroblasts in the insert for 6 or 10 days, respectively, prior to microbial inoculation. The apical surface of the epithelial layer was inoculated with optimal (three strains of *L. crispatus*) or non-optimal (two strains of *G. vaginalis* and one strain of *P. amnii*) reconstructed consortia. After 48 hours, both consortia showed stable growth on inserts containing A2EN or HCxEC cells (**Figure 3A**). No significant differences were observed in the relative growth of the individual strains within the optimal consortium on a HCxEC epithelium over 48 hours, indicating viability and growth from all three strains (**Figure 3B**). Similarly, all members of the non-optimal consortium grew efficiently over the 48-hour co-culture period, without significant variation on growth rates (**Figures 3AC**). Viability staining demonstrated that the bacteria localized in close proximity to the host epithelial surface (**Figure 3D**). A live/dead assay confirmed that neither the optimal nor the non-optimal consortia induce cytotoxicity in HCxEC cervical cells (**Figure 3E**).

**Fig. 3.**
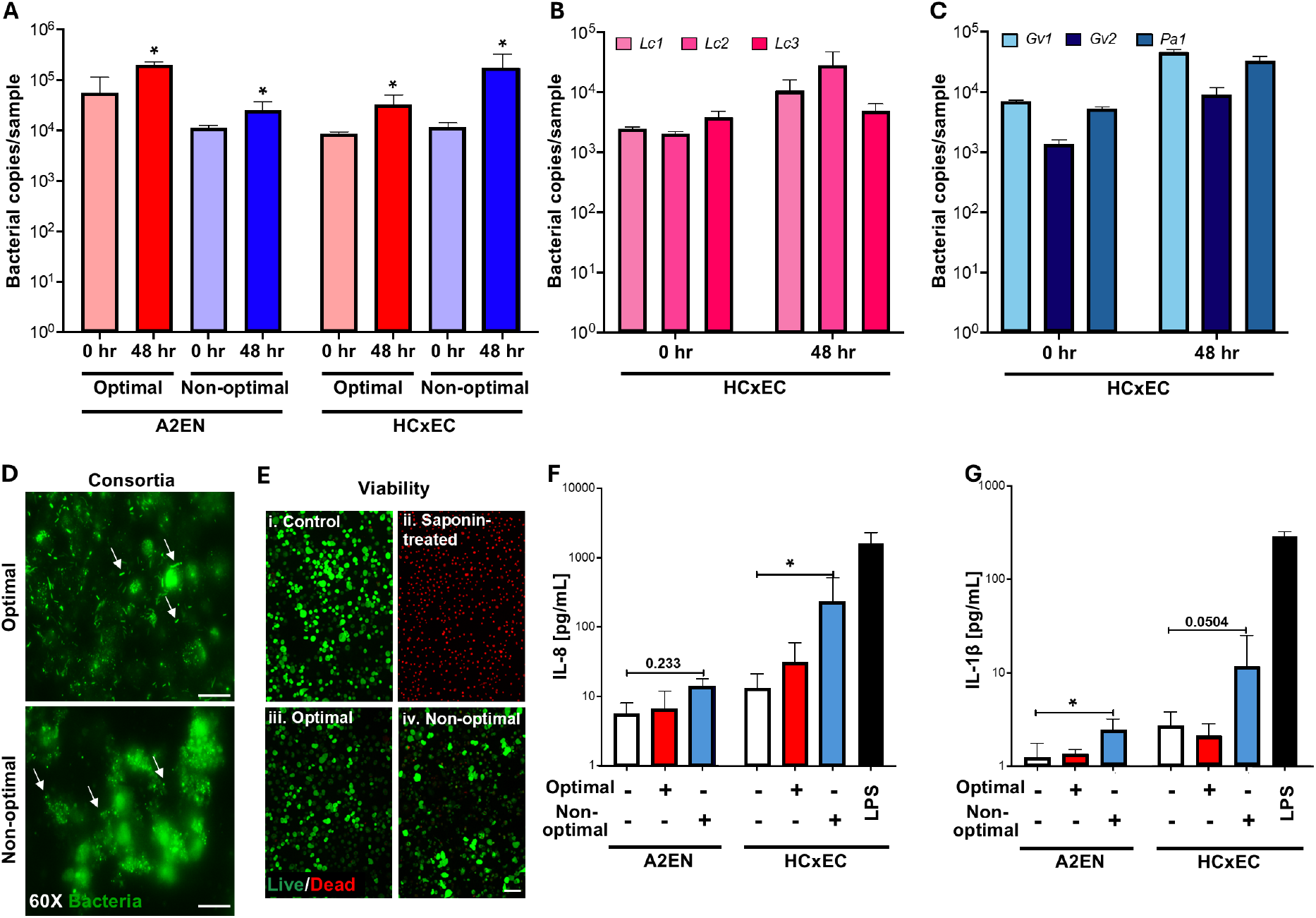
Microbiota consortia grow in the MPS device and do not elicit negative epithelial response. **(A)** Growth of the optimal and non-optimal microbiota consortia after 48 hours in the insert on A2EN and primary cervical (HCxEC) cells. Stability of the growth of optimal **(B)** and non-optimal **(C)** microbiota consortia in the insert on primary cervical cells (HCxEC) after 48 hours. **(D)** Bacteria comprising the optimal and non-optimal microbiota consortia (arrows) are alive after culture on epithelium. Scale bar, 10 µm. **(E)** Live/dead viability staining of HCxEC cells after 48 hours of growth without (i,ii) or with (iii-iv) each consortium and HCxEC cells treated with saponin (ii). Scale bar, 20 µm. **(F)** IL-8 and **(G)** IL-1β cytokine response of A2EN and HCxEC cells in the basal channel of the insert 48 hours after co-culture with optimal and non-optimal microbiota consortia. Significance determined by ANOVA * = p<0.05

To investigate host responses, we quantified secretion of pro-inflammatory cytokines IL-8 and IL-1*β* in the basal channel following 48-hour co-culture of A2EN or HCxEC cells with either the optimal or non-optimal consortia or no bacteria. At baseline, cytokine levels were approximately two-fold higher in HCxEC cells compared to A2EN cells (**Figure 3FG**), consistent with the previously reported truncated immune response of the A2EN cell line(42). For both epithelial models, exposure to the optimal consortium induced negligible changes in IL-8 and IL-1*β* levels (**Figure 3FG**). In contrast, exposure to the non-optimal consortium elicited distinct pro-inflammatory responses. A2EN cells exhibited a modest two-fold increase in IL-8 and IL-1*β* levels, whereas HCxEC cells showed an 18-fold increase in IL-8 levels and a 4-fold increase in IL-1*β* levels (**Figures 3FG**). Together, these findings demonstrate that cervical epithelial cells can be stably co-cultured with defined reconstructed microbiota consortia within the MPS device and that, as *in vivo*, host inflammatory responses differ depending on the associated microbiota(58). These results validate the physiological relevance of the MPS device and highlight its utility for dissecting how complex microbial communities influence epithelial inflammation and susceptibility to infection, an advance over previous monoassociation systems(17, 59).

### *Ct* infects and completes a full developmental cycle

Once internalized into susceptible host cells, *Ct* undergoes a biphasic developmental cycle within a membranebound compartment called the inclusion, whereby infecting elementary bodies (EBs) differentiate into replicating reticulate bodies (RBs) soon after internalization. RBs differentiate “back” to infectious EBs late in development in preparation for host cell lysis and reinfection. To determine if *Ct* could successfully complete this developmental cycle in the MPS device, we infected A2EN cells co-cultured with BJ fibroblasts. The infection and development were monitored with a strain of *Ct* that constitutively expresses mCherry by visualizing inclusions in the epithelial compartment at different times post-infection (**Figures 4AB**). Inclusions were observed throughout the A2EN monolayer, indicating successful and homogeneous infection across the reconstituted epithelium (**Figure 4A**). Inclusions increased in size from 24 to 48 hours post-infection (hpi), consistent with *Ct* growth and inclusion expansion. By 72 hpi, inclusion lysis was evident, indicating progression to the final stage of the *Ct* developmental cycle (**Figure 4B**). To further confirm the bacteria developmental cycle progression through the asynchronous transition of RBs to EBs starting in mid-cycle (from 20 to 48+ hpi for *Ct*), we infected A2EN cells co-cultured with BJ fibroblasts with a *Ct* reporter strain in which mCherry is constitutively expressed and GFP is expressed at the beginning of the RB-to-EB transition and in mature EBs (**Figure 4C**). At 24 hpi, the majority of the *Ct* inclusions were mCherry-positive but GFP-negative, consistent with the predominance of RBs. By 48 hpi, inclusions were positive for both mCherry and GFP, indicating transition to the EB stage (**Figure 4C**) and successful progression through the developmental cycle. Quantification of inclusion size using mCherry (RBs; **Figure 4D**) and GFP (EBs; **Figure 4D**) confirmed both inclusion growth and RB-to-EB transition from 24 to 48 hpi. To further demonstrate the successful completion of the *Ct* developmental cycle and validate the MPS as a physiologically relevant model of *Ct* infection, we showed that infectious EBs could be recovered from the insert in an MOIdependent manner (**Figure 4E**). This directly indicates that the MPS device supports productive *Ct* infection, with the ability to yield new, infectious EBs capable of initiating subsequent rounds of infection.

**Fig. 4.**
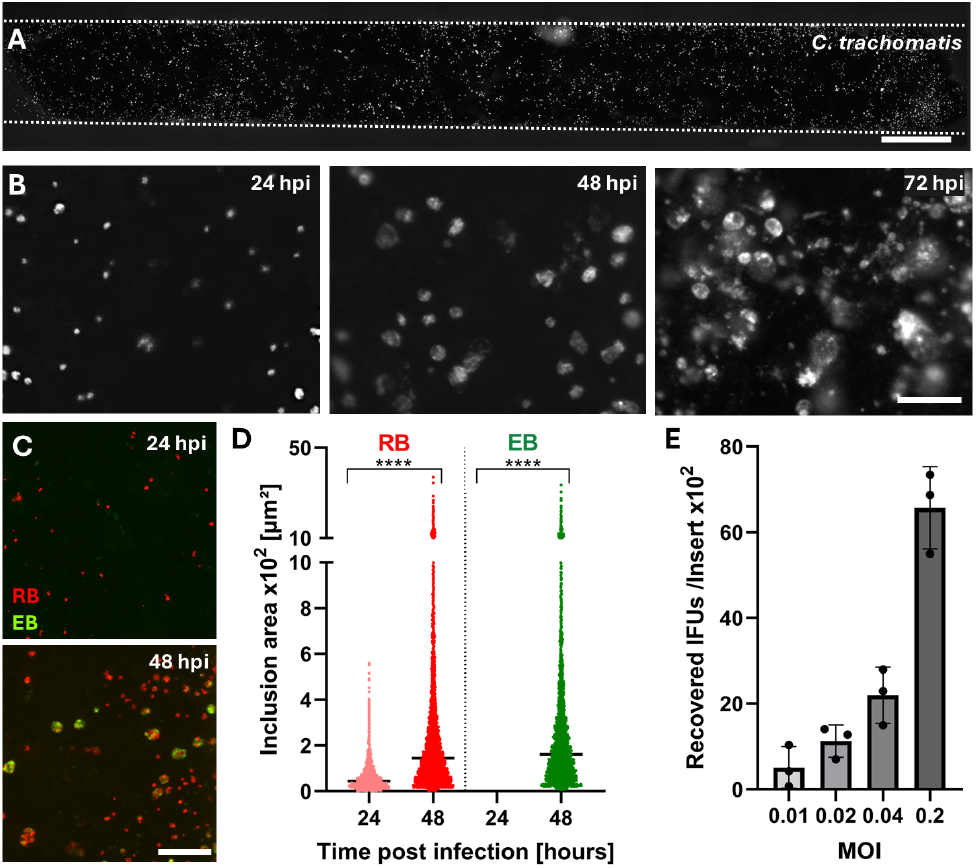
*C. trachomatis* undergoes a complete developmental cycle in A2EN cervical cells co-cultured with BJ fibroblasts in a MPS device. **(A-B)** Epifluorescence microscope images of the epithelial channel cultured under static conditions and infected with mCherry-expressing *Ct*. The mCherry channel is shown; each white dot represents *Ct* inclusions. **(A)** Low magnification of the entire channel at 48 hpi. Dotted lines indicate channel edge. Scale bar, 1 mm. **(B)** Representative images at higher magnification at 24 (left), 48 (middle), and 72 (right) hpi. Scale bar, 100 µm. **(C)** Epifluorescence microscopy merged images of A2EN epithelial cells in the apical channel of the MPS device, cultured under static conditions and infected with *Ct* expressing mCherry constitutively (red) and GFP under the control of a late developmental cycle promoter (green). Scale bar, 100 µm. **(D)** Quantification of the size of RB and EB containing inclusions at 24 and 48 hpi from three independent experiments. Mann-Whitney tests, **** p < 0.0001. **(E)** Quantification of recovered IFUs per insert at 48 hpi in response to increasing inoculum concentrations. Data represent three independent experiments and are presented as mean ± SEM.

### Pre-exposure to optimal microbiota reduces *Ct* infectivity and secondary propagation

To assess the effects of the reconstructed microbiota consortia against *Ct* infection, primary cervical HCxEC cells were cultured in the MPS insert, and on day 10, cocultured with either the optimal or non-optimal microbiota consortium (2×10^4^ bacteria/insert). After 48 hours, *Ct* was added at an MOI of 1, and the number of infected cells was measured 48 hours later. Cells co-cultured with the non-optimal microbiota consortium exhibited similar levels of infection (30%) as untreated controls (**Figure 5AB**). In contrast, co-cultures with the optimal microbiota consortium reduced *Ct* infection by more than 3-fold (8%) (**Figure 5AB**). To determine whether co-culture with the optimal microbiota consortium also limited secondary propagation, *Ct* viability and infectiousness, lysates from primary infections of HCxEC cells were used to infect HeLa cell monolayers. At 24 hpi, a 6-fold reduction in secondary infection was observed compared to the control (**Figure 5CD**). These findings show that co-culture of primary cervical cells with the optimal microbiota consortium not only reduces primary *Ct* infection but also impairs the infectivity of progeny EBs produced by the initial infection. Together, these results highlight that chlamydial infection and propagation and the protective capacity of optimal microbiomes to reduce chlamydial infection can be replicated in primary cervical cells cultured in the MPS device.

**Fig. 5.**
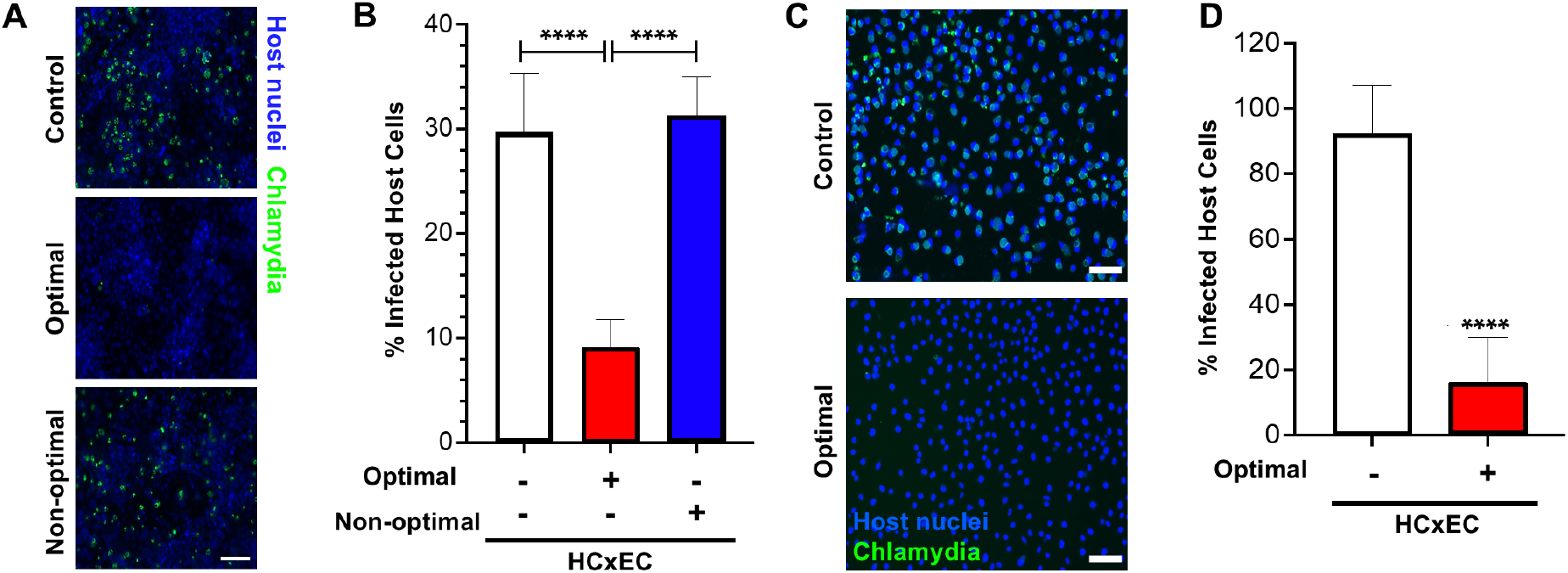
Protective capacity of the optimal microbiota consortium on HCxEC cells from *Ct* infection. **(A)** Representative images of HCxEC cells co-cultured with BJ fibroblasts after 48 hours of co-culture with the optimal or non-optimal microbiota consortium. HCxEC nuclei are stained in blue, and chlamydiae are stained in green. Scale bar, 20 µm. **(B)** Quantification of infection with and without microbiota based on the number of cells with chlamydiae. **(C)** Representative images of secondary chlamydial infection in HeLa cells after HCxEC culture with or without microbiota. HeLa cell nuclei are stained in blue, and chlamydiae are stained in green. Scale bar, 20 µm. **(D)** Quantification of secondary infection in HeLa cells based on the number of cells with chlamydiae. **** p<0.0001

### *Ng* infects and replicates on cervical epithelia under static and flow conditions

*Ng*, a facultative intracellular pathogen, can replicate on the apical surface of an epithelium or within the epithelial cells or neutrophils(32). To test *Ng* infection in the MPS model, the apical surface of A2EN cells co-cultured in the insert with BJ fibroblasts was inoculated with *Ng* (2×10^6^ CFU) under static conditions for 1 hour. At 8 hpi, microscopy and colony forming units (CFU) counts indicated robust *Ng* colonization and replication, similar to *Ng* growth in the MPS device that only contained culture medium (**Figure 6AB**). To evaluate *Ng* infection under flow, inserts containing A2EN cells co-cultured with BJ fibroblasts were clamped in the flow cassette and exposed to 2.5 µL/min flow. After 6 hours, *Ng* remained adherent to the epithelium despite continuous flow, confirming tight interaction with the A2EN cells (**Figure 6C**). The assessment of *Ng* growth on the surface of the epithelial cells is confounded by the fact that the observed bacteria represent a population that reflects replication, recruitment from other locations, and loss from being washed away. To clarify whether the observed *Ng* were replicating within the apical epithelial channel a dual-labeling approach was used. GFP-expressing *Ng* prelabelled with CTFR lose the CTFR signal as they replicate while maintaining the constitutively produced GFP signal. By 6 hrs. post infection 48% of the *Ng* were GFP positive and CTFR negative, suggesting the observed bacterial population had undergone replication within the MPS. (**Figure 6C**). These findings confirm that the MPS system supports not only *Ng* adherence *per se*, but also flow-stable adherence and colonization, a critical aspect of *Ng* mucosal infection not captured in static culture models.

**Fig. 6.**
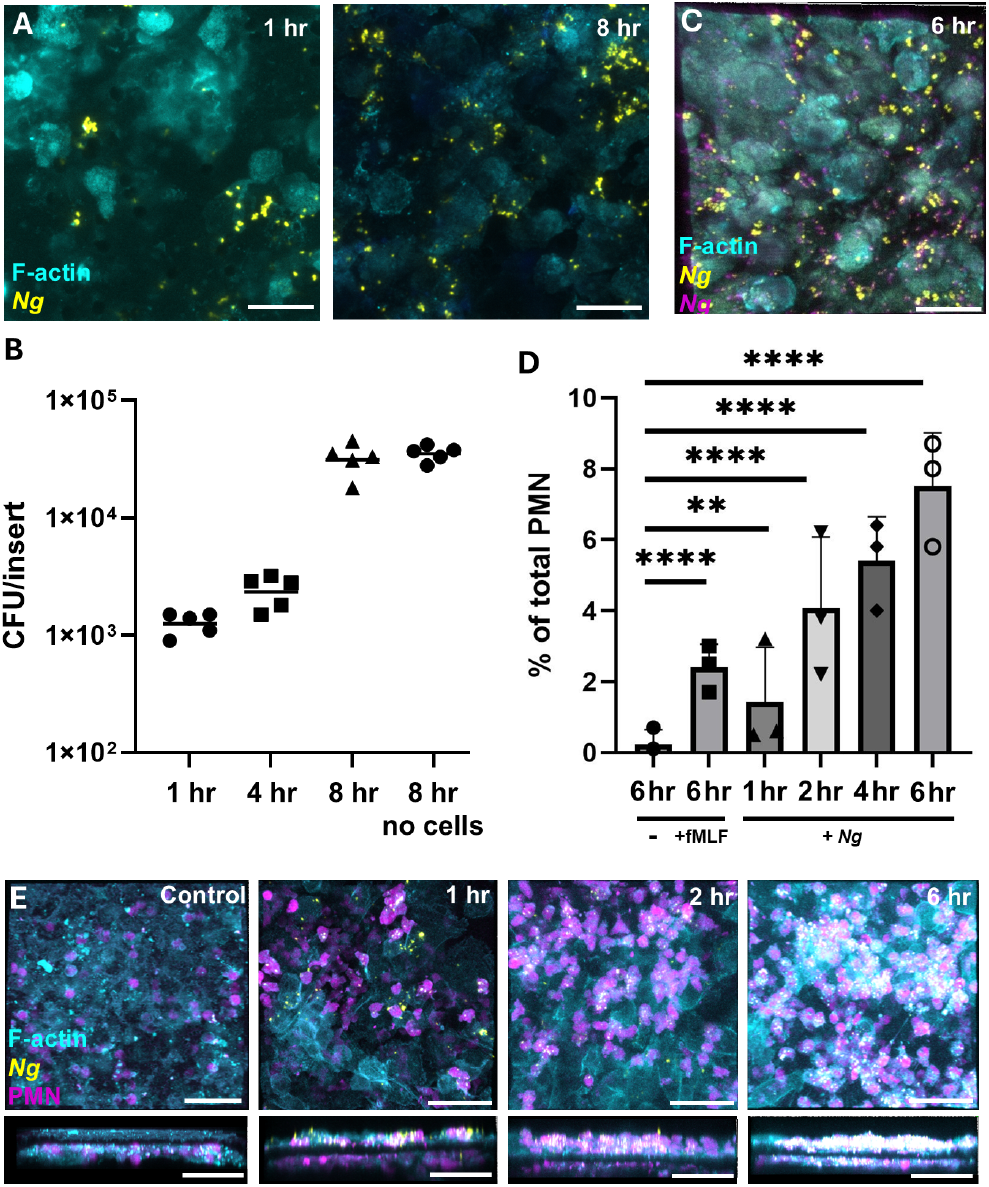
*Ng* infection of A2EN cells and consequent neutrophil transepithelial migration in the MPS device. **(A)** *Ng* infection (yellow) at 1 hour and 8 hours post inoculation on A2EN and BJ fibroblast (cyan) laden inserts. Scale bar, 20 µm. **(B)** Quantification of *Ng* growth over time. Scale bar, 20 µm. Growth of *Ng* in cell culture medium in the MPS device without A2EN/fibroblast cells (no cells) is presented for comparison. **(C)** Infection of A2EN cells (cyan) with GFP+ (yellow) *Ng* that was prelabeled with Cell Trace Far Red (magenta). Cells were incubated for 6 hours under 2.5 µL/min flow conditions. As *Ng* replicates, magenta fluorescence is lost. **(D)** Quantification and **(E)** visualization of neutrophil (pink) migration through the insert in response to *Ng* (yellow) infection of the cervical epithelium (cyan), (timelapse movie: **Supplemental Video S1**). *Ng* was added for at t=-1 hr, PMN were added at t =0, images and measurements were taken at t = 1, 2, 4, 6 hr. Positive control, inserts to which the PMN chemoattractant fMLF was added for 6 hours; negative control, no fMLF addition and no *Ng*. Scale bar, 40 µm.

### Neutrophils transmigrate across the cervical epithelium in response to *Ng* infection

Local neutrophil recruitment is a hallmark of gonococcal infection *in vivo*(60). To recapitulate this host immune response to *Ng* infection in the MPS device, we co-cultured A2EN cells with BJ fibroblasts on inserts, infected the epithelial layer with *Ng* for 1 hour, and added PMNs to the basal channel in contact with the BJ fibroblasts to mimic neutrophils in circulation. fMLF in the epithelial channel was used as a positive control chemoattractant, while uninfected, untreated inserts were a negative control for PMN recruitment. PMN transmigration into the epithelial channel in response to *Ng* significantly increased over 6 hours, compared to uninfected controls (**Figure 6DE, Supplemental Video S1**). Thus, the MPS model reproduces a key host immune response to infection, i.e., immune cell recruitment to the infected site, and can be used to study epithelial-immune-pathogen interactions during gonococcal pathogenesis.

## Discussion

The objective of the work was to cooperatively develop a cost-effective, transferable MPS device that recapitulates the basic physiology and cellular architecture of cervical tissue to investigate host-pathogen interactions in a biomimetic environment. Specifically, we focused on two prominent sexually transmitted pathogens, *Chlamydia trachomatis* (*Ct*) and *Neisseria gonorrhoeae* (*Ng*). In addition to supporting infection, the device was designed to reproduce key host defense mechanistic elements, including mechanical (e.g., fluid flow), microbial (i.e., optimal and non-optimal microbiota), and immune (e.g., neutrophil recruitment and inflammatory response) components. Here, we present a comprehensive characterization of an MPS device of the cervical epithelium that fulfills these requirements.

Using A2EN endocervical epithelial cells and HCxEC primary cervical cells co-cultured with BJ fibroblasts, we engineered an MPS device that reproduces tissue-like morphology and barrier functions, features consistent with cervicovaginal physiology. The device enables controlled fluid flow through the epithelial and basal channels, creating shear stresses known to play a role in proper cell physiology(61). While fluid flow control is a feature of all microfluidic devices, our design enabled non-engineering users to establish flow simply and easily, creating a device that is more accessible than standard microfluidic devices.

To enhance the model’s biomimicry and relevance for studying STI pathogenesis, we co-cultured defined optimal and non-optimal microbiota with the cervical epithelium in the MPS device. Despite their known clinical relevance, little is known about the mechanisms by which vaginal microbiota modulate susceptibility to infection. Epidemiological studies have consistently shown that women with vaginal microbiota lacking Lactobacillus spp. and enriched in strict and facultative anaerobes, characteristic of bacterial vaginosis, are at significantly increased risks for *Ct* and *Ng* infection after exposure to an infected partner(4, 62–65). Thus, these microbiota are non-optimal or ‘permissive’ by virtue of their association with an increased risk of infection. Conversely, Lactobacillus-dominated vaginal microbiota are associated with reduced susceptibility to infections. Studying the mechanisms by which these microbiota are protective or permissive requires the development of improved *in vitro* models that mimic the cervicovaginal microenvironment, including cervical mucus, immune components, and a persistently colonizing cervicovaginal microbiota. Most current *in vitro* models for studying *Ct* and *Ng* pathogenesis often rely on infecting immortalized cervical cells and lack the host response primed by optimal or non-optimal cervicovaginal microbiota that is present before infections, resulting in an incomplete characterization of host-pathogen interactions.

Attempts to recapitulate host-microbiota interactions have used simple *in vitro* co-culture systems consisting of immortalized vaginal or cervical human epithelial cells exposed to single species or strains of vaginal bacteria(66). These systems do not faithfully replicate important functions of the cervicovaginal space, as they lack differentiated epithelium and an interstitial compartment. In recent studies, vaginal epithelial cells were differentiated into polarized multilayers and exposed to air and bacteria(67). These co-culture models revealed a modest anti-inflammatory effect of *L. crispatus* and *L. jensenii* after innate immune stimulation(67). We previously addressed some of these limitations by developing a 3D organotypic cervical and vaginal epithelial models with collagen-embedded fibroblasts, which supported single-strain microbial growth(17). However, these models, although relatively inexpensive, reproducible, and experimentally tractable, still lack the full functional capacities of the cervicovaginal microenvironment, including a persistently colonizing cervicovaginal microbiota. For these reasons, these models have major limitations for studies of STI pathogenesis.

A key advance of the current model is the successful co-culture of defined, simplified optimal (multiple strains of *Lactobacillus crispatus*) and non-optimal (*Gardnerella vaginalis* and *Prevotella amnii*) vaginal microbiota with differentiated cervical epithelial cells supported by BJ fibroblasts, in the MPS device. This system allowed for sustained microbial viability, spatial proximity to host cells, and differential modulation of cytokine responses, without compromising epithelial barrier integrity. These features enable mechanistic interrogation of how specific microbiota influence epithelial health, inflammation, and susceptibility to infection. These advances underscore the MPS model’s ability to recapitulate the clinically observed dichotomy between protective and permissive microbiota states. The device enables the controlled manipulation of microbial composition, providing a human-relevant platform for directly testing causal relationships between microbiota composition and host outcomes, which is not achievable in static cultures or animal models. This capability is a significant advancement over traditional models, which typically lack the stability, spatial resolution, or immune integration to support such studies.

To further support the utility of the platform as a promising tool for advancing our understanding of microbiota-driven susceptibility to infection, we demonstrated the impact of optimal and non-optimal vaginal microbiota on the outcomes of *Ct* infections. We demonstrated that *Ct* could successfully infect the reconstituted epithelium and undergo its full developmental cycle in the device. Fluorescence imaging confirmed formation, expansion, and transition of *Ct* inclusions from RBs to EBs, validating the MPS platform as a physiologically relevant model of *Ct* infection. As expected, HCxEC cells co-cultured with the non-optimal microbiota consortia showed *Ct* infection rates comparable to those of uncolonized controls. In contrast, cervical cells associated with the optimal microbiota showed significantly reduced *Ct* infection, in line with epidemiological data. These findings indicate that the MPS not only supports the intracellular development of *Ct* but also faithfully recapitulates the protective effects of *L. crispatus*-dominated microbiota. Importantly, pre-colonization of the primary cervical epithelium with non-optimal microbiota in the MPS enabled infection in a more physiologically relevant context, one in which the host epithelium is primed by its resident microbiota prior to pathogen exposure.

While the simplified three-member consortia used here captured key microbiota-level functions, future studies may expand microbial complexity to include additional bacterial species and strains associated with non-optimal and permissive microbiota, explore temporal dynamics of microbiota succession and transition, or incorporate pH modulation or metabolite gradients to further enhance physiological relevance.

Using the MPS device we also tested *Ng* infection. *Ng* adhered to and proliferated on A2EN-based endocervical epithelia under both still and flow conditions, features consistent with *Ng* natural mucosal colonization and infection. Single cell-type, static 2D tissue culture assays have revealed that *Ng* uses an array of surface structures to promote adherence to human epithelial cells, including type IV pili, Opa proteins, lipooligosaccharide, and the porin PorB(32). The MPS platform allows us to define the relative contribution of these and other surface features to endocervical infection under flow compared with static conditions. Indeed, surface adhesion by uropathogenic *Escherichia coli, Staphylococcus aureus*, and *Pseudomonas aeruginosa*, which uses type IV pili similarly to *Ng*, are enhanced by shear forces(68–70), demonstrating the need to account for fluid flow when modeling sites of bacterial infection with cell culture. Relatedly, epithelial cell expression profiles are significantly altered when exposed to shear stress(71); whether similar changes affect cellular binding partners for *Ng* adhesins, and resulting consequences for *Ng* infection, remain to be determined.

Upon *Ng* infection, PMNs added to the basal channel of the MPS device migrated through the fibroblast layer, the membrane, and the A2EN epithelium to the apical surface, where they engulfed gonococci. These findings recapitulate the PMN influx seen in mucosal human *Ng* infection(72). Previously, we demonstrated that *Ng*-infected endocervical cells seeded on permeable filter supports enable the basal-to-apical recruitment of human PMNs by the coordinate release of two lipid chemoattractants(16). Epithelial cells produce hepoxilin A3 when in intimate contact with *Ng*, and the resulting recruited PMNs secrete leuktotriene B4, amplifying the PMN response76. The MPS presented here allows us to test the contribution of these chemoattractants and other signaling pathways in a model that is more complex given the architecture and the presence of subepithelial fibroblasts. Though often considered only in the context of extracellular matrix dynamics, fibroblasts can both upregulate or suppress the immune response, depending on the context(73). The MPS affords the opportunity to dissect this signaling and integrate additional immune components found in the endocervix, including resident innate and adaptive immune cells, to faithfully capture the complete dynamics of infection with *Ng* and other STI pathogens(72). One unanswered question in the *Ng* field is the outcome of PMN recruitment for the bacteria and the epithelium. With the MPS, future studies can define how differences in *Ng* adhesin expression and binding of opsonins (complement, IgG) affect bacterial phagocytosis by PMNs that have epithelially transmigrated. Since the antimicrobial proteins, peptides, and reactive oxygen species made by PMNs can also cause tissue damage(74), future experiments with the MPS can also reveal how ongoing PMN influx in response to infection affects *Ng* survival as well as the viability and integrity of the PMNs and epithelium.

One of our major objectives was to democratize access to physiologically relevant MPS models of the cervicovaginal tract that mimic human tissues, enabling advanced investigations of STI host-pathogen interactions that are not possible using current model systems. Microfluidic models of the cervix have been previously characterized(36), however, these use specialized equipment and complex techniques that make them inaccessible to non-engineering users. Our team of bioengineers and researchers cooperatively designed an insert-cassette MPS device that was modular, low-cost, and compatible with standard laboratory workflows, requiring no cleanroom facilities. Importantly, we demonstrated successful technology transfer from bioengineering researchers to multiple independent laboratories, overcoming a key challenge in organon-chip systems: reproducibility. By developing the device in one lab and successfully translating materials and protocols to multiple independent labs, the cervical MPS device has demonstrated reproducibility. These results support broader implementation of this platform beyond the founding laboratories and potentially beyond the study of chlamydial and gonococcal infections.

Our MPS device also offers distinct advantages over traditional models. Unlike static Transwell® cultures, it enables dynamic flow and immune cell transmigration. Compared to organoid systems, it allows for easier access by microbes and the immune system. And unlike animal models, it offers human tissue specificity, including its unique microbiota, and experimental controls. The platform is adaptable for future extensions, including hormonal cycling, menstrual blood simulation, and co-infection studies (e.g., *Ct–Ng* or *Trichomonas vaginalis*). Nevertheless, we acknowledge that the current MPS system supports relatively short-term experiments and limited microbiota diversity and complexity. Additionally, while neutrophil migration assays were reproducible and robust, they were limited to acute responses over several hours. Longitudinal experiments, scaling for higher throughput, and use of clinical isolates will be critical next steps for translation and refinement.

In summary, we have successfully developed a microfluidic device that faithfully mimics the cervical microenvironment, supports the complete infection cycles of *Ct* and *Ng*, and responds appropriately to key host-specific modulators such as vaginal microbiota and innate immune cells. This platform lays the groundwork for translational applications, including screening of live biotherapeutics, evaluation of microbicides, and studies of host-microbiome interactions. More broadly, the MPS device offers a novel democratized, easy-to-use, and affordable model suitable for advancing our understanding of STI pathophysiology and accelerating the development and testing of novel therapeutics.

## ACKNOWLEDGEMENTS

This work was supported in part by a grant from the National Institutes of Health (U19AI158930).

## DATA AND RESOURCE AVAILABILITY

Requests for further information and resources, see https://www.gleghornlab.com/resources.

## AUTHOR CONTRIBUTIONS

Conceptualization (KMN, DJM, VLE, IJG, DJDD, KT, FCW, PMB, ID, AKC, JR, JPG) Formal analysis (KMN, DJM, VLE, IJG, DJDD) Funding acquisition (PMB, ID, AKC, JR, JPG) Investigation (KMN, DJM, VLE, IJG, DJDD, KT, FCW) Methodology (KMN, DJM, VLE, IJG, DJDD, KT, FCW) Project administration (KMN, PMB, ID, AKC, JR, JPG) Supervision (PMB, ID, AKC, JR, JPG) Validation (KMN, DJM, VLE, IJG, DJDD, KT, FCW) Visualization (KMN, VLE, IJG, DJDD) Writing-original draft (KMN, VLE, IJG, DJDD) Writing – review & editing (KMN, DJM, VLE, IJG, DJDD, KT, FCW, PMB, ID, AKC, JR, JPG)

## Supplementary Information

**Fig. S1.**
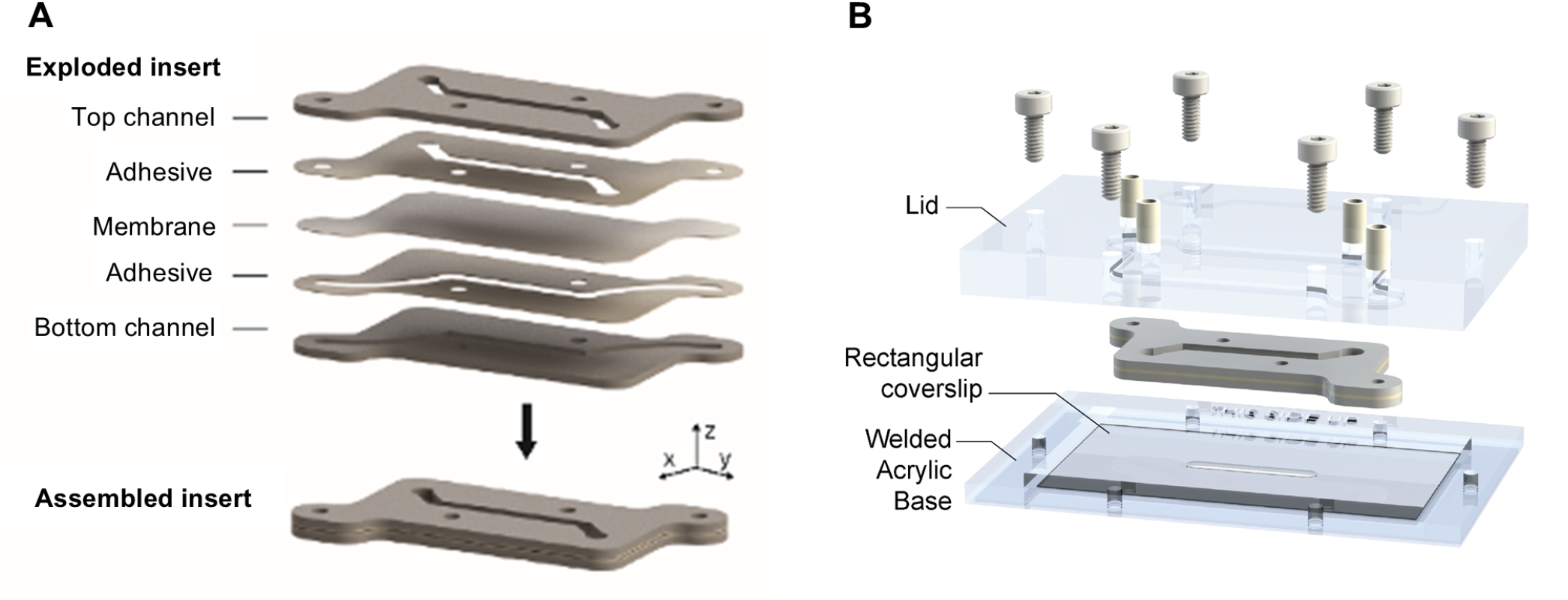
Design of the insert and the cassette. **(A)** An exploded view of the insert showing the different layers that are used to assemble the insert. **(B)** The insert gets clamped into an acrylic cassette to enable fluid flow and live imaging with an inverted microscope.

**Fig. S2.**
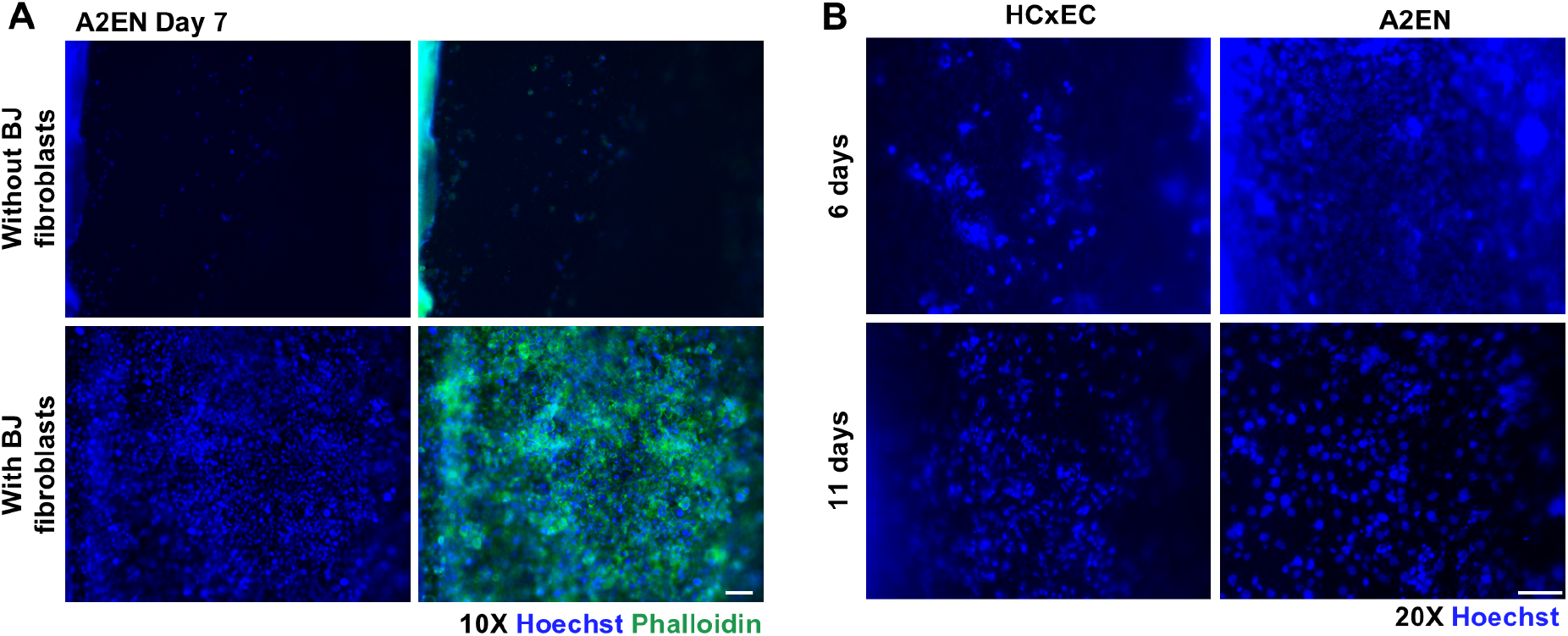
Requirements for consistent microphysiological growth of cervical cells. **(A)** A2EN cervical cells require the addition of BJ fibroblasts feeder cells to facilitate confluent growth. Nuclei are stained with Hoechst (blue) and actin is stained with phalloidin (green). Merged channels are shown in the right column. **(B)** Primary (HCxEC) and A2EN cervical cells attain confluency at different time points. Scale bars, 20 µm.

**Table S1.**
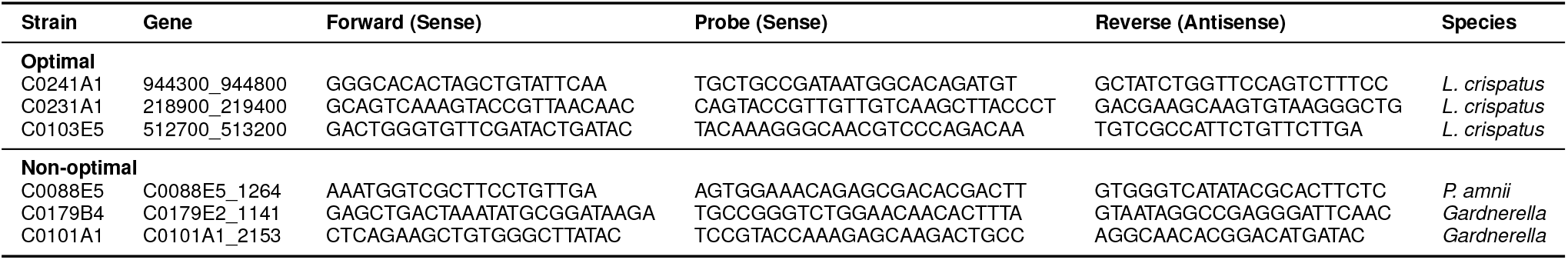
Primer and probe sequences for each bacterial strain used.

**Video S1.**
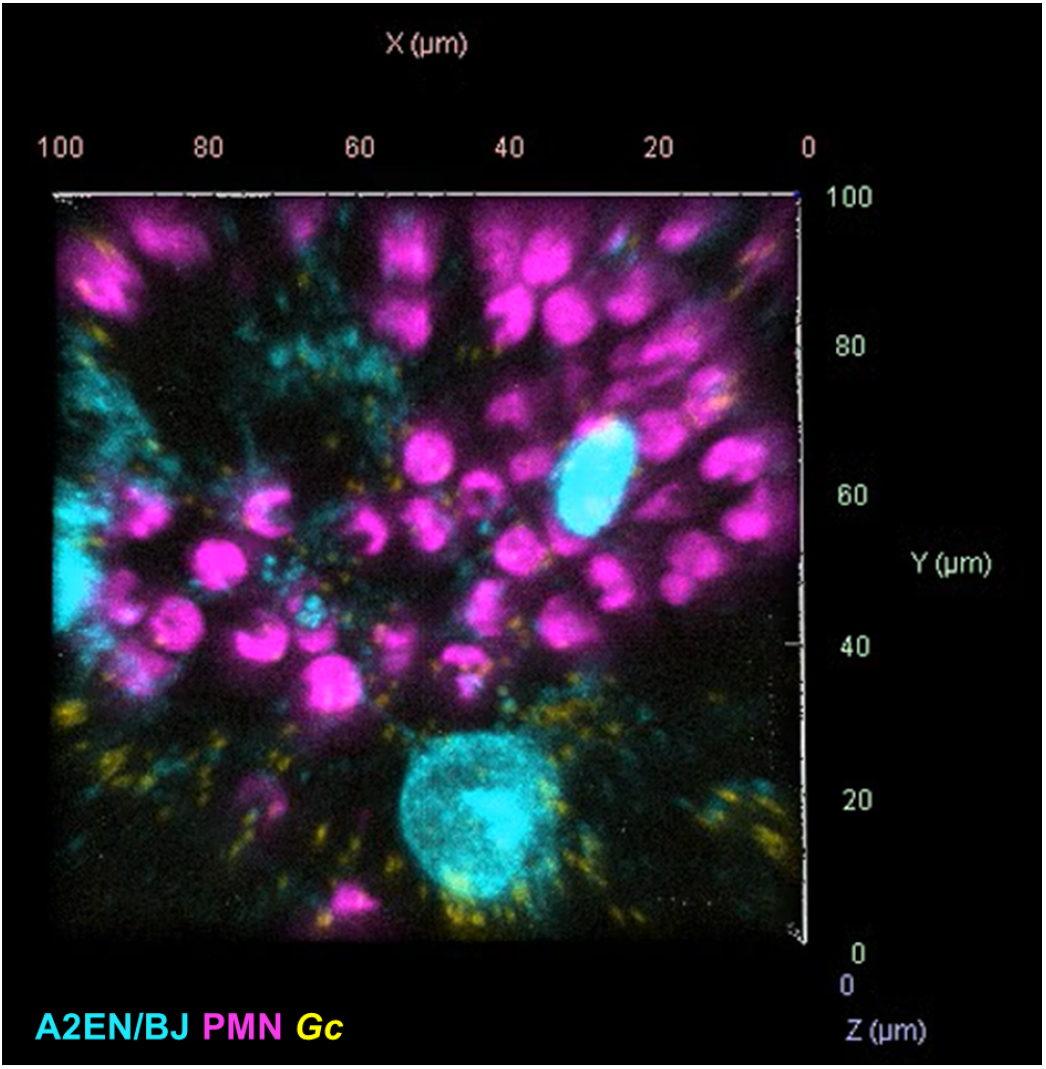
Timelapse video of PMN migration from the basal channel to the apical surface of an A2EN epithelium. Yellow= GFP bacteria, Cyan= Cell Trace Yellow A2EN and BJ cells, Magenta= CellTrace Far Red PMN. Video taken from 2-3 hours post PMN addition. Video can be found at https://gleghornlab.com/resources.

